# Soft selective sweeps in evolutionary rescue

**DOI:** 10.1101/052993

**Authors:** Benjamin A. Wilson, Pleuni S. Pennings, Dmitri A. Petrov

**Affiliations:** Department of Biology, Stanford University, Stanford, CA 94305 USA; Department of Biology, San Francisco State University, San Francisco, CA 94132 USA

**Author notes:** **Corresponding author:** Ben Wilson, Department of Biology, Stanford University, 371 Serra Mall, Stanford, CA, 94305.

## Abstract

Evolutionary rescue occurs when a population that is declining in size because of an environmental change is rescued by genetic adaptation. Evolutionary rescue is an important phenomenon at the intersection of ecology and population genetics. While most population genetic models of evolutionary rescue focus on estimating the probability of rescue, we focus on whether one or more adaptive lineages contribute to evolutionary rescue. We find that when evolutionary rescue is likely, it is often driven by soft selective sweeps where multiple adaptive mutations spread through the population simultaneously. We give full analytic results for the probability of evolutionary rescue and the probability that evolutionary rescue occurs via soft selective sweeps in our model. We expect that these results will find utility in understanding the genetic signatures associated with various evolutionary rescue scenarios in large populations, such as the evolution of drug resistance in viral, bacterial, or eukaryotic pathogens.

---

Abrupt environmental changes may lead to a demographic decline in a population when the population is maladapted to the new environment. This scenario necessitates an evolutionary response from the population if the population is to escape extinction. The process by which genetic adaptation allows a population to recover from the demographic consequences of harsh environmental shifts has been termed “evolutionary rescue”. Evolutionary rescue is a focus of many studies at the interface of ecology and evolution in part due to recent attention to climate change, drug resistance, pesticide resistance, and other anthropogenic change of global and local environments (Gonzalez *et al.* 2013; Alexander *et al.* 2014; Carlson *et al.* 2014). While much attention has been given to theoretical and experimental predictions regarding the probability that evolutionary rescue occurs under various scenarios, we focus on the dynamics of adaptive alleles and the associated genetic signatures left behind by adaptation. More explicitly we are interested in how selective sweeps that enable evolutionary rescue differ in the number of ancestral backgrounds on which they emerge, based on the conditions under which evolutionary rescue occurs. If evolutionary rescue is facilitated by ‘hard’ selective sweeps—wherein a single progenitor is responsible for the spread of the beneficial variant—then genetic diversity will be removed from the population as a result of adaptation as well as demographic decline. By contrast, a population that adapts via ‘soft’ selective sweeps—wherein multiple ancestors have independently derived beneficial variants— will preserve some of the ancestral diversity that was present prior to the environmental shift that caused the population to decline (Pennings and Hermisson 2006b).

Soft selective sweeps occur when adaptive alleles appear in multiple individuals prior to sweeping through the population. This leads to a sample genealogy that includes multiple adapted ancestors in the new environment. The underlying criterium for the occurrence of soft selective sweeps is that the presence of adaptive mutations in the population is not a limiting factor to the process of adaptation (Hermisson and Pennings 2005; Pennings and Hermisson 2006a; Burke 2012; Messer and Petrov 2013). This criterium is fulfilled in many situations, such as adaptation from previously neutral or deleterious standing genetic variation (Hermisson and Pennings 2005), adaptation from recurrent *de novo* mutation (Pennings and Hermisson 2006a), or adaptation in a spatially structured metapopulation (Ralph and Coop 2010). While soft selective sweeps appear to be abundant in case studies of adaptation (Messer and Petrov 2013), the signature of a soft selective sweep is crucially dependent on the population sample and the underlying demographic history of the population from which the sample is taken. Importantly, many case studies of adaptation pertain to situations where the demography of the population and the process of adaptation are necessarily interrelated. Particularly in the case of resistance evolution, beneficial mutations of large effect often confer benefits in terms of absolute fitness rather than relative fitness, thereby leading to demographic changes to the population. Previous work suggests that soft selective sweeps in populations with fluctuating population size can give rise to signatures of both hard and soft sweeps depending on when beneficial alleles arise, when they are sampled, and how advantageous they are (Wilson *et al.* 2014). However, in that work we had assumed that demography was independent of the allelic state at the locus under selection, an assumption that is valid only under models where fitness advantages are relative and density-independent (*e.g.* the standard Wright-Fisher model with selection or the Moran model). In this paper, we allow adaptation to influence demography. We explore a simple logistic model where fitness advantages are absolute and density-dependent within the context of an evolutionary rescue scenario. While most models of evolutionary rescue predict the probability that rescue occurs, we focus on the likelihood that rescue occurs via hard or soft sweeps.

The primary result of our analysis is that when rescue is likely to occur, it is more likely to occur via soft selective sweeps than hard selective sweeps. This result follows intuitively from the observation that if the time-averaged input of adaptive mutations is higher, then this will lead to a greater probability of rescue and a greater probability of observing soft selective sweeps. We demonstrate how our result is critically dependent on the population-scale mutation rate at the onset of the wildtype population decline and whether mutant growth rates are restricted by population density. We give analytical results for the probability of evolutionary rescue and the probability that rescue occurs via soft selective sweeps. We also give the waiting-time distributions for rescuing beneficial mutations. In the context of previous results connecting soft sweeps and post-adaptation genetic diversity, our results highlight a key correlation between genetic diversity following evolutionary rescue and the likelihood of evolutionary rescue (Feder *et al.* 2016).

## Results

We begin by modeling a population that is going extinct because of an environmental shift that leads to a demographic decline in the population (*e.g.* a drug enters the host environment of a virus causing the viral population to crash). We assume that mutants are not present at the onset of the population decline, *i.e.* there are no adaptive mutations present as standing genetic variation. Adaptive mutations emerge on the background of the maladapted wildtype. We assume a single-locus, two-allele model where the mutation rate toward the beneficial state is *μ* and is constant in time. We assume back mutations to the wildtype state are negligible. Individuals of the two types, maladapted wildtype (*w*) and adapted mutant (*m*), give birth or die with transition rates given by

- *w* → *w* + 1: *b*_*w*_*w*
- *w* → *w* - 1: *d*_*w*_*w*
- *m* → *m* + 1: 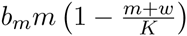
- *m* → *m* - 1: *d*_*m*_*m*.

The decline of the wildtype (maladapted) population is intrinsic to the genotype and density-independent, *i.e.* the wildtype suffers decreased reproductive success directly from its interaction with the environment and not from competition for shared resources. The wildtype population size can be deterministically approximated by *w*(*t*) = *w*_0_exp(-*αt*) where the variable *α* sets the rate at which the wildtype population declines and is equivalent to the absolute difference in per capita birth and death rates (*α* = |*b*_*w*_ – *d*_*w*_|). The carrying capacity *K* sets the scale of density dependence and determines the equilibrium population size for the adapted population should adaptation occur: *m*_eq_. = *K* (1 – *d*_*m*_/*b*_*m*_), which is the value of *m* obtained by setting the mutant birth rate 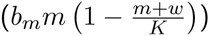 equal to the mutant death rate (*d*_*m*_*m*), setting *w* = 0, and solving for *m*.

For simplicity we will assume that *d*_*m*_ = *d*_*w*_ such that the generation time required for the life cycle of each type is the same. In an alternative model, this assumption could be relaxed to investigate how genetic alterations to generation time or how different reproductive strategies (such as viral latency) could facilitate evolutionary rescue, but we will not explore these scenarios in this investigation. Ignoring the density-dependent scaling factor 1 – (*m* + *w*)/*K*, the difference *b*_*m*_ – *b*_*w*_ could be interpreted as the genotype-intrinsic growth advantage of a mutant individual over a wildtype individual. The parameter *b*_*m*_ also sets the maximum per capita birth rate for mutants at low population density. For our model, we do not consider extremely large birth rates (*b*_*m*_ ⪢ 1) to avoid extreme jumps in the population size over short timescales. Note that this model is closely analogous to scenario 2 under *Alternative Forms of Population Regulation* in Orr and Unckless (2008).

At any given time *t*, a mutation has a probability of appearing, *w*(*t*)*μ*, and a probability of establishing, *p*_est_.(*t*). Establishment occurs when a mutation survives extinction due to drift at low copy number. From analysis of soft sweeps via *de novo* mutation in populations of constant size, we know that soft sweeps are only expected to occur when *w*_0_*μ* ~ 1 or greater (Pennings and Hermisson 2006a). In most scenarios we consider, we correspondingly scale *μ* to be either 1/*w*_0_ or 10/*w*_0_ to ensure that mutations occur frequently enough at the beginning of the environmental shift to expect multiple adaptive lineages to appear during the rescue scenario, though their survival will ultimately depend on the other parameters previously described, namely the carrying capacity (*K*), the wildtype decline rate (*α*), and the mutant birth rate (*b*_*m*_). Note that the decline in the wildtype population means that adaptation will eventually be mutation-limited in all cases. We also consider situations where adaptation is mutation-limited (*w*_0_*μ* ⪡ 1) to illustrate the limitations of our model assumptions. Figure 1 gives an illustration of the rescue scenario.

**Figure 1:**
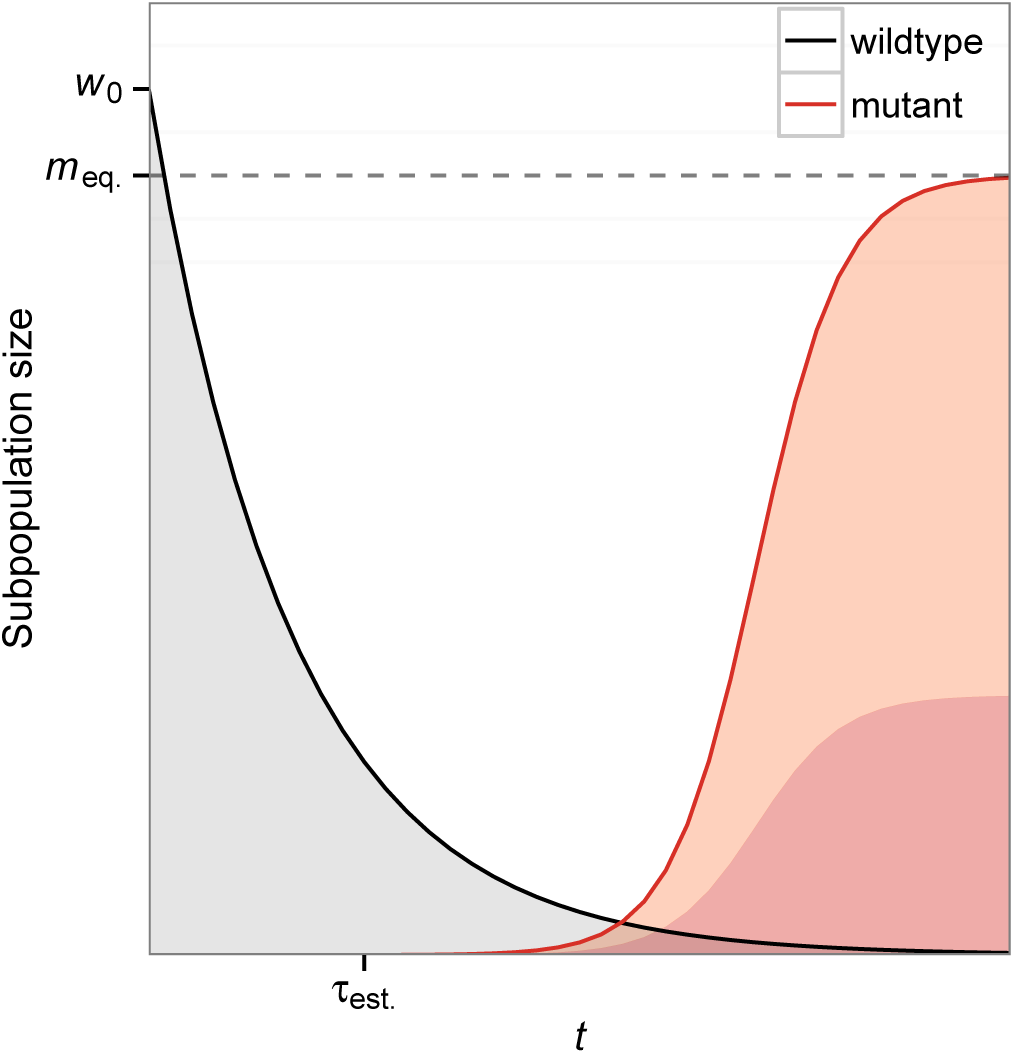
Model depiction of evolutionary rescue. A population initially composed of maladapted wildtype individuals (black) declines exponentially from its original size *w*_0_ following an environmental shift. A beneficial mutation appears on the background of a wildtype individual and establishes at time *τ*_est_., at which point the population is destined to be rescued via adaptation. If a mutant fails to establish before the wildtype population goes extinct, then no rescue occurs. Following rescue, the mutant population will equilibrate at a new population size m_eq_. = *K* (1 – *d*_*m*_/*b*_*m*_). In some cases of rescue, multiple mutant lineages can establish before the wildtype population goes extinct, leading to a soft selective sweep as illustrated by the multiple shaded lineages (red) within the mutant subpopulation.

### The time-dependent probability of establishment

We can derive the probability of establishment for a mutation arising at a particular time *τ* using the methodology presented in Uecker and Hermisson (2011) (see specifically equation A5 under *Fixation in General Ecological Models* in the appendix and equation 16a in the main text). Uecker and Hermisson (2011) showed that for a time-inhomogeneous birth-death process (such as the specific birth-death model presented here) we can write the probability of establishment for a single mutant starting at particular time *τ* = 0 in terms of the total per capita mutant birth rate *B*(*t*) and total per capita mutant death rate *D*(*t*). The general result takes the form

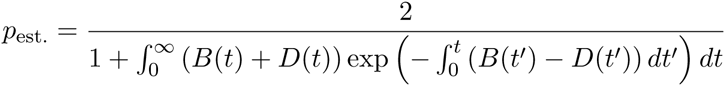

where *t*′ is a dummy variable for the nested integral. For our specific birth-death model, 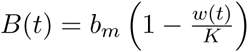 and *D*(*t*) = *d*_*m*_ can be taken directly from the transition rates presented at the beginning of this section, and the probability of establishment for a single mutant appearing at a particular time *τ* is given by

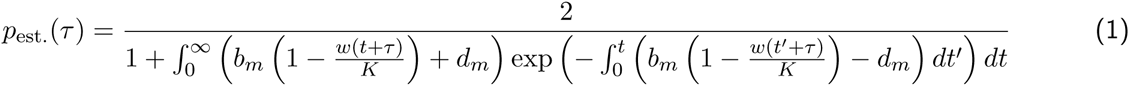

where the instantaneous time *τ* now appears explicitly because *τ* is not fixed at zero.

Note that we have neglected the mutant population size in the density-dependent terms under the assumption that mutant lineages have independent probabilities of establishment while the mutant population size is low and while the expected time between successive establishments is short. Later we will show that this assumption breaks down for rescue scenarios with slow decline rates when the population-scale mutation rate is low.

### The role of population density

We demonstrate here how population density influences the process of evolutionary rescue in our model. Including density dependence through a carrying capacity *K* ensures that a rescued population reaches an equilibrium size at long timescales. Population density also has critical effects on mutant establishment because it determines the growth rate of the mutant through time. Note that population density here is the population size relative to the carrying capacity, not the population size itself.

We can separate the effects of population density into two characteristics: the growth rate of the mutant at the onset of the wildtype population decline and the rate at which density-dependent growth restriction decays over time. The initial growth rate of the mutant depends on the ratio of the starting wildtype population size to the total population size limit (*w*_0_/*K*). If the initial population density is high, *i.e.* the carrying capacity is smaller than or similar in size as the wildtype population size, then the birth rate of the mutant may (initially) be lower than its death rate, making it unlikely that a mutant establishes. This is the case when the following inequality holds: *w*_0_/*K* > 1 - *d*_*m*_/*b*_*m*_.

By contrast, in situations where the carrying capacity is much larger than the wildtype population size, a mutant lineage that appears at the onset of the wildtype population decline will have a net positive growth rate from the onset. This holds when *w*_0_/*K* < 1 - *d*_*m*_/*b*_*m*_. In other words, early mutants have almost no chance of survival when population density is initially high but are able to establish when population density is initially low.

The rate of population decline of the wildtype population determines how quickly mutant growth rates increase. Scenarios with fast wildtype population decline will alleviate mutant growth restrictions more quickly and increase the probability of establishment (conditional on appearance), although fast wildtype population decline will also decrease the rate at which mutants appear. We highlight these different aspects of density dependence in our model because we find it important to note that 1) early mutants are not unconditionally advantageous compared to wildtype individuals, and 2) the probability of establishment is intrinsically tied to the wildtype population decline.

The influence of population density on *p*_est_.(*t*) can be seen by comparing values of *p*_est_.(*t*) between situations of high and low population density at the onset of the wildtype population decline, as illustrated in Figure 2A. In the scenario with high population density (*w*_0_/*K* > 1-*d*_*m*_/*b*_*m*_), *p*_est_.(*t*) is essentially zero until the wildtype population has declined sufficiently to give mutants a positive growth rate (bottom, red line). This growth rate transition occurs near *w*\* ~ *K*(1-*d*_*m*_/*b*_*m*_). Using the deterministic approximation to the wildtype population size, we can estimate that this transition occurs at

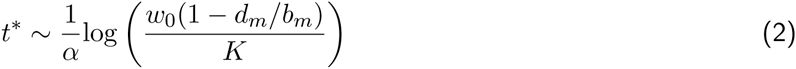

by setting *w*(*t*) = *w*\* = *K*(1 - *d*_*m*_/*b*_*m*_), and solving for *t*. By contrast, the scenario with low population density at the onset (*w*_0_/*K* < 1 - *d*_*m*_/*b*_*m*_) shows that beneficial mutants have an appreciable probability of establishing from the beginning of the environmental shift (top, blue line). Note that because the probability of establishment is measured for a mutation occurring at time *t* and because the density restriction imposed by the wildtype population declines monotonically with time, *p*_est_. will increase monotonically with time in our model. It asymptotes at a value

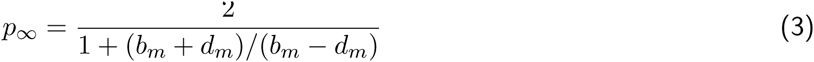

that is obtained by taking the limit as *t* → ∞ for Equation (1) and is therefore independent of *K* and *α* because the wildtype population will eventually go extinct.

**Figure 2:**
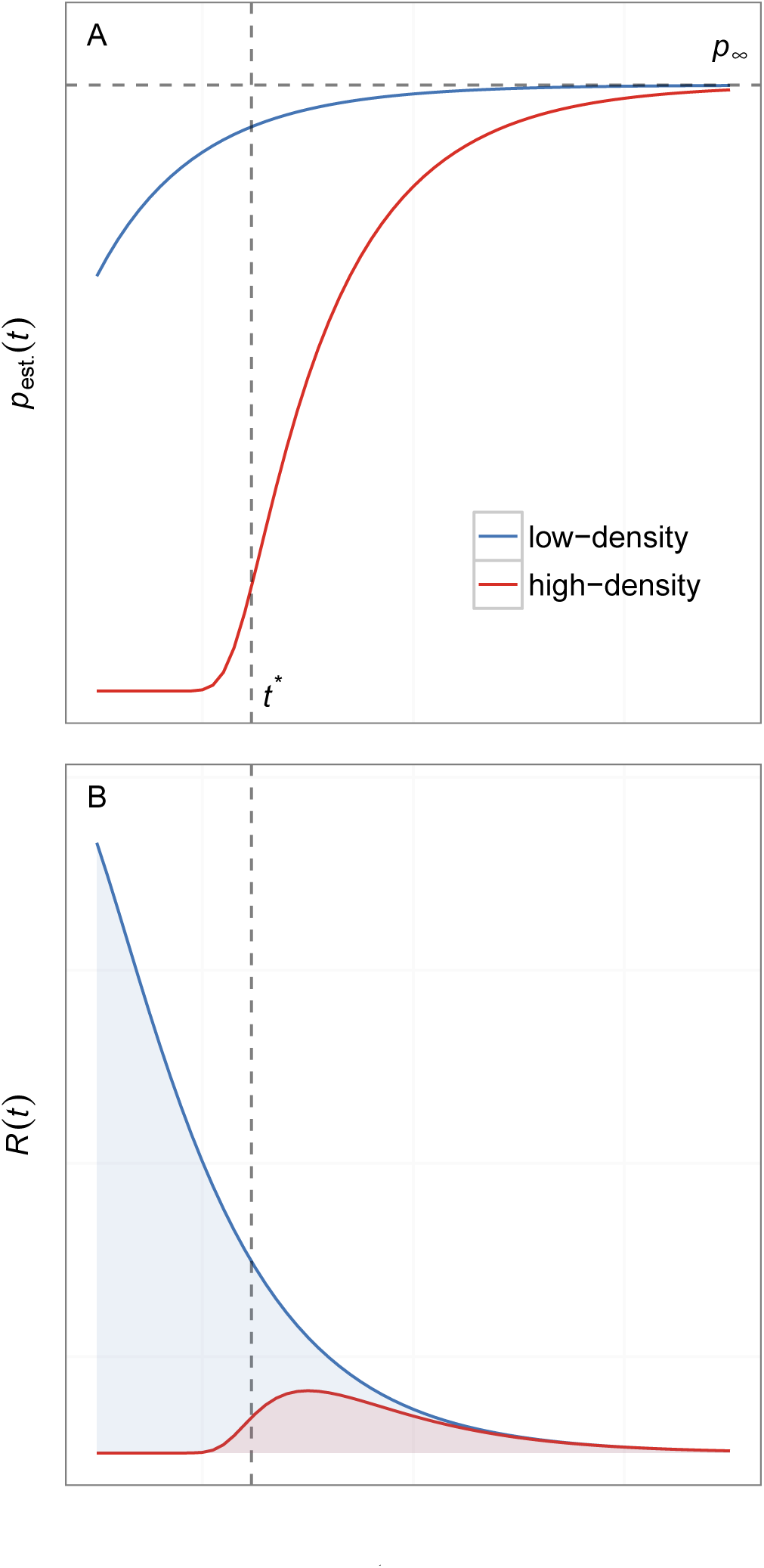
Establishment probability and intensity function distributions. Part A shows the establishment probability distributions for mutations appearing at time *t* in low-(top, blue) and high-density (bottom, red) scenarios with all parameters being equal except for *K*. For low-density scenarios, the probability of establishment increases monotonically as the density restriction on mutant growth declines to zero with the wildtype population. For high-density scenarios, the probability of establishment is essentially zero until the wildtype population declines to a size *w*\* at *t*\* (see Equation (2)) whereupon it will increase monotonically just as in the low-density scenario (these approximations can be conservative). Note that both low-and high-density scenarios have distributions that asymptote to the same value, p_∞_, because p_∞_ is only dependent on the unscaled per capita birth and death rates of the mutant (see Equation (3)). Part B shows the distributions for the corresponding intensity functions *R*(*t*) = *w*(*t*)*μp*_est_.(*t*) for the same two scenarios in Part A. *R*(*t*) gives the instantaneous rate at which mutants successfully establish and save the population from extinction. *R*(*t*) eventually declines to zero with the wildtype population size (see Figure 1) even though the establishment probability increases with time. The shaded area under *R*(*t*) determines the probability of evolutionary rescue and is generally larger in low-density (top, blue) scenarios than in high-density scenarios (bottom, red).

Although the probability of establishment increases over time, the rate of appearance of mutants (*w*(*t*)*μ*) decreases monotonically coinciding with the wildtype population size decline. Thus at long times, the total rate of successful beneficial mutants, *R*(*t*) = *w*(*t*)*μp*_est_.(*t*), will eventually decay with time as shown in Figure 2B). In the following section, we show how *R*(*t*) can be used as the intensity function for a time-inhomogeneous Poisson process that determines mutant establishments.

### Evolutionary rescue by soft selective sweeps

With these model considerations in mind, we can derive the probability that a population headed for extinction is rescued by at least one successful adaptive mutant. The probability of rescue is 1 – *P*_extinction_, or one minus the probability that no mutants establish. The probability that no mutants establish is exp 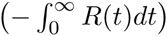 if we model mutant establishments using a time-inhomogeneous Poisson process with intensity function *R*(*t*) = *w*(*t*)*μp*_est_.(*t*). This leads to a total probability of rescue equal to

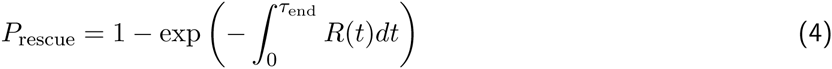

where we have replaced the upper limit ∞ with *τ*_end_ = log(*w*_0_)/*α* representing the time it would take for a deterministically declining wildtype population to reach a single individual. The integral in Equation (4) is the area under the intensity function depicted in Figure 2B and represents the number of mutants expected to establish during the time when mutations can occur in wildtype individuals.

Assuming independence between mutant lineages, we can gain an overall picture of whether rescue is more likely to occur via hard or soft selective sweeps using the same time-inhomogeneous Poisson process to model the establishment of each individual lineage. To determine the probability of evolutionary rescue via soft selective sweeps, we will first want to calculate that the probability that only one mutant establishes before the wildtype population goes extinct, *i.e.* evolutionary rescue occurs via a hard selective sweep. This is given by

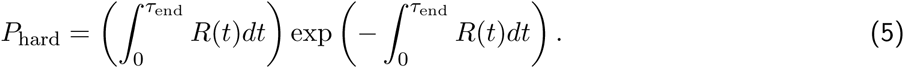

Evolutionary rescue requires at least one mutant lineage to establish before the wildtype population goes extinct, and all evolutionary rescue that does not occur via a hard sweep must occur via a soft sweep by definition. Therefore, the probability of evolutionary rescue via soft selective sweeps is

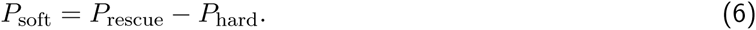

To confirm our analysis, we performed forward-time birth-death simulations in populations of 10, 000 individuals over multiple values of *K*, *α*, *b*_*m*_, and *μ* (see Methods section for details). If we examine *P*_rescue_ for low-density (*w*_0_ = 10,000, *K* = 110,000) and high-density ( *w*_0_ = 10,000, *K* = 10,000) rescue scenarios, we can see that the overall probabilities of rescue and rescue via soft sweeps decline with increasing *α* (see simulation and analytic values from Equation (4) and Equation (6) plotted in Figure 3 and Figure 4). The qualitative dependence on decline rate in our model is the same as seen in previous models where mutation was weak (Orr and Unckless 2008, 2014). As for the other relevant parameters in our model, the probabilities of evolutionary rescue and rescue via soft sweeps increase universally with increasing *μ* and *b*_*m*_. Rescue is generally higher in low-density scenarios (when the carrying capacity is higher than the wildtype population size) than in high-density scenarios (when the carrying capacity is close to the wildtype population size), similar to simulation results for scenario 2 under *Alternative Forms of Population Regulation* in Orr and Unckless (2008). We also find that sweeps are generally softer in low-density scenarios than in high-density scenarios.

**Figure 3:**
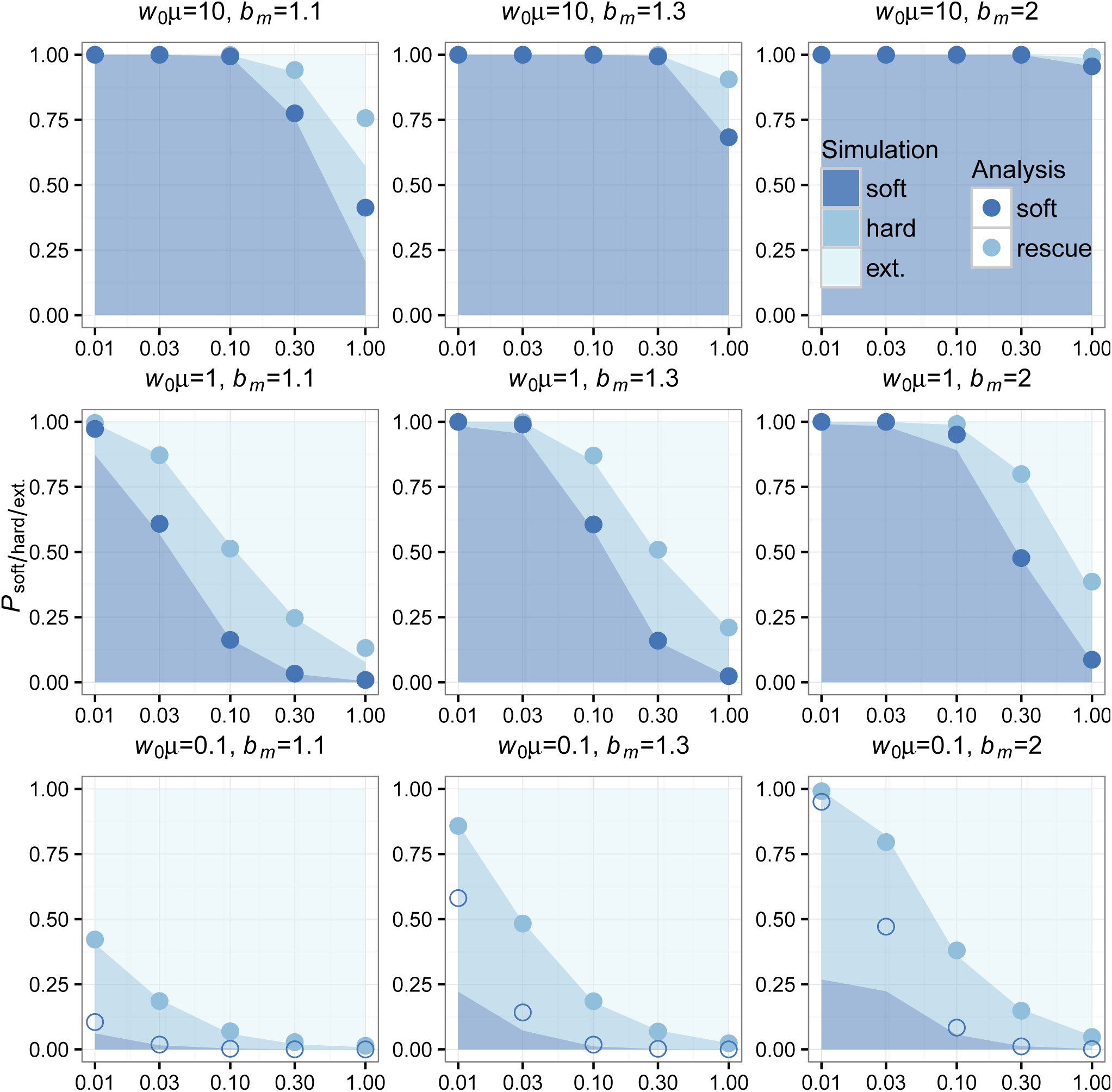
Simulations and analytic predictions for evolutionary rescue in low-density scenarios. The probabilities of observing rescue via soft selective sweeps, rescue via hard selective sweeps, or extinction as a function of the decline rate *α* (logarithmic scale) measured over 1000 simulations (see Methods) for each combination of model parameters are indicated as cumulative bar plots. The color key indicates shading for simulation outcomes. Population-scale mutation rate increases between plots from bottom to top, and unscaled per capita mutant birth rate increases between plots from left to right. The analytic predictions for each parameter combination show *P*_soft_ (bottom blue points) and *P*_rescue_ = *P*_hard_ + *P*_soft_ (top, light blue points). For analytic predictions of *P*_soft_, points where adaptation is not mutation-limited are shaded (*w*_0_*μ* ≥ 1). Our analysis has high concordance with the observed probability of rescue for each parameter combination. Our analysis also has high concordance with the observed probability of rescue via soft sweeps for most parameter combinations, except in instances where evolutionary rescue is mutation-limited (*w*_0_*μ* ⪡ 1) from the onset for low decline rates (bottom row, leftmost *α* values).

**Figure 4:**
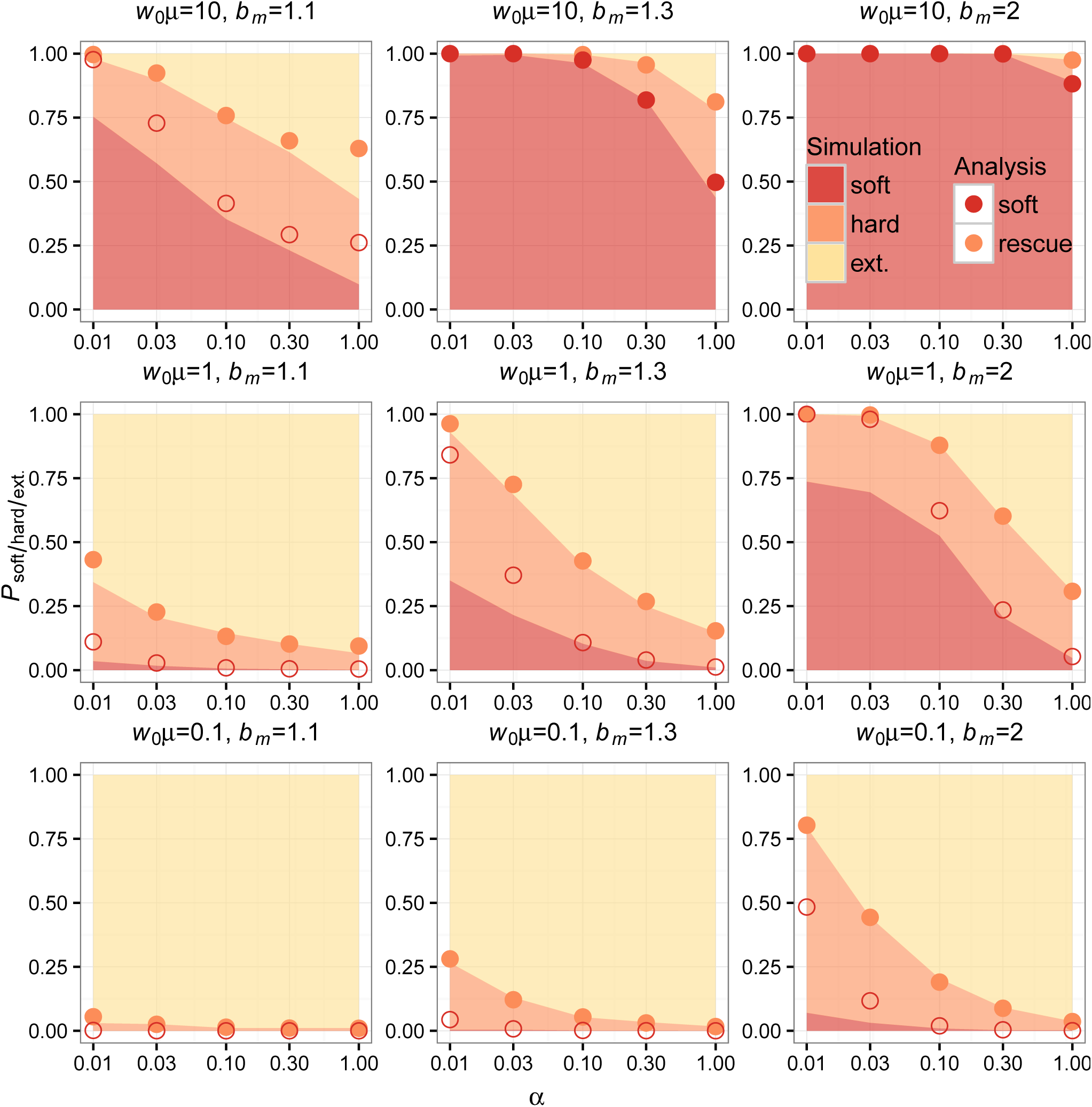
Simulations and analytic predictions for evolutionary rescue in high-density scenarios. The probabilities of observing rescue via soft selective sweeps, rescue via hard selective sweeps, or extinction as a function of the decline rate *α* (logarithmic scale) measured over 1000 simulations (see Methods) for each combination of model parameters are indicated as cumulative bar plots. The color key indicates shading for simulation outcomes. Population-scale mutation rate increases between plots from bottom to top, and unscaled per capita mutant birth rate increases between plots from left to right. The analytic predictions for each parameter combination show *P*_soft_ (bottom red points) and *P*_rescue_ = *P*_hard_ + *P*_soft_ (top orange points). For analytic predictions of *P*_soft_, points where adaptation is not mutation-limited are shaded (*w*\**μ* ≥ 1). Our analysis has high concordance with the observed probability of rescue for each parameter combination. As in the low-density scenario, our analysis has good concordance with the observed probability of rescue via soft sweeps for most parameter combinations, except in instances where evolutionary rescue is mutation-limited (*w*\**μ* ⪡ 1) and for low decline rates (bottom and middle rows, leftmost *α* values). Evolutionary rescue is mutation-limited for more parameter combinations in the high-density scenario because the wildtype population must decline before establishment of mutants is likely (see Figure 2), making the initial population-scale mutation rate effectively lower and the area under *R*(*t*) smaller.

Simulations and analysis agree well for both low-and high-density scenarios when comparing corresponding values of *P*_rescue_. Our analysis also agrees with the observed probability of rescue via soft sweeps in scenarios where adaptation is not limited by mutation (see Figure 3 and Figure 4). In instances where evolutionary rescue is mutation-limited (*w*_0_*μ* < 1 for low-density rescue and *w*\**μ* < 1 for high-density rescue) and where the wildtype population declines slowly, our analytic assumption regarding the independence of mutant lineages during establishment breaks down resulting in deviations from the values observed in simulations. In particular, our analysis overestimates the probability of soft sweeps because it excludes the contribution of the mutant subpopulation in the density-dependent term in Equation (1).

### Mutation-limited evolutionary rescue

Although the primary focus of this paper is to investigate soft selective sweeps in evolutionary rescue, we feel that mutation-limited evolutionary rescue deserves special attention. When evolutionary rescue is mutation-limited, the population will typically either go extinct or be rescued by a single mutant lineage via a hard selective sweep. Neither of these outcomes should impact the validity of our assumption of independence between lineages in Equation (1) because they involve either zero or one mutant lineages respectively. However, in mutation-limited scenarios where the wildtype population declines very slowly and where the time between mutant establishments is long, it is possible that one mutant lineage establishes and reaches a size large enough to prevent a second mutant lineage from establishing, leading to a hard sweep. This scenario is not accounted for in our analysis because we exclude the mutant contribution to population density in the density-dependent term in Equation (1). In these situations the window of opportunity for a second mutant lineage to establish is limited by the time it takes for the first established mutant lineage to bring the total population size high enough to substantially decrease the establishment probability of a second mutant lineage, assuming the wildtype population declines slowly enough to allow multiple mutations to appear before it goes extinct. This is similar to the scenario in Pennings and Hermisson (2006a) where a second mutant establishing was limited by the time it takes the first mutant to sweep in a population. What is most noteworthy about these situations is that our observations of whether evolutionary rescue appears mutation-limited or not depend on population density at the onset of the environmental shift, not just the population-scale mutation rate at the onset of the environmental shift. Scenarios where one established mutant lineage can prevent others from establishing occur more frequently in the high-density scenarios than in the low-density scenarios that we explored. This is because in high-density scenarios, mutant lineages are unlikely to establish until the wildtype population declines to *w*\* ~ *K*(1 – *d*_*m*_/*b*_*m*_), at which point the population-scale mutation rate can often be limiting (*w*\* < 1) to adaptation even when *w*_0_*μ* ≥ 1. Understanding when this phenomenon will occur and when we expect evolutionary rescue to appear mutation-limited is therefore dependent on approximating the population-scale mutation rate when mutant lineages have an appreciable probability of establishing. We have distinguished scenarios where mutant establishment probabilities are approximately independent (*w*\**μ* ≥ 1) from scenarios where our assumptions regarding independence between lineages are expected to break down because evolutionary rescue is mutation-limited in Figure 3 and Figure 4.

### Waiting-time distributions for the establishment of mutants

It may be of interest to reformulate the results from our previous analysis in terms of the waiting times associated with the establishment of the individual adaptive lineages during the extinction process. As in the previous analysis, we will assume independence between mutant lineages. Under this assumption, we define *τ*_1_ to be the waiting time for the first mutant lineage to establish. *τ*_1_ has probability density equal to

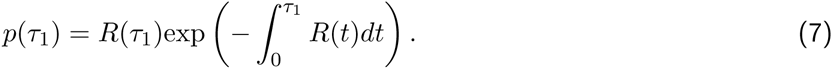

Conditional on the establishment of a first adaptive mutant at *τ*_1_, the probability density for the establishment of a second adaptive mutant at time *τ*_2_ takes the same form integrated over all possible *τ*_1_. The probability density for *τ*_2_ is

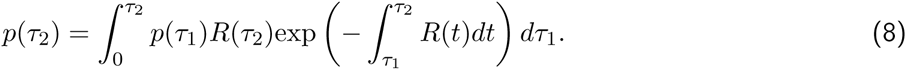

Distributions for both *τ*_1_ and *τ*_2_ are plotted for one set of parameters in Figure 5. Equation (7) and Equation (8) are in good agreement with forward-time birth-death simulations for this parameter combination, although it is important to consider the previous discussion regarding the independence of mutant lineages during establishment and the regime where independence breaks down, in which case we expect departures for the distribution for *τ*_2_ but the distribution for *τ*_1_ should remain unchanged.

**Figure 5:**
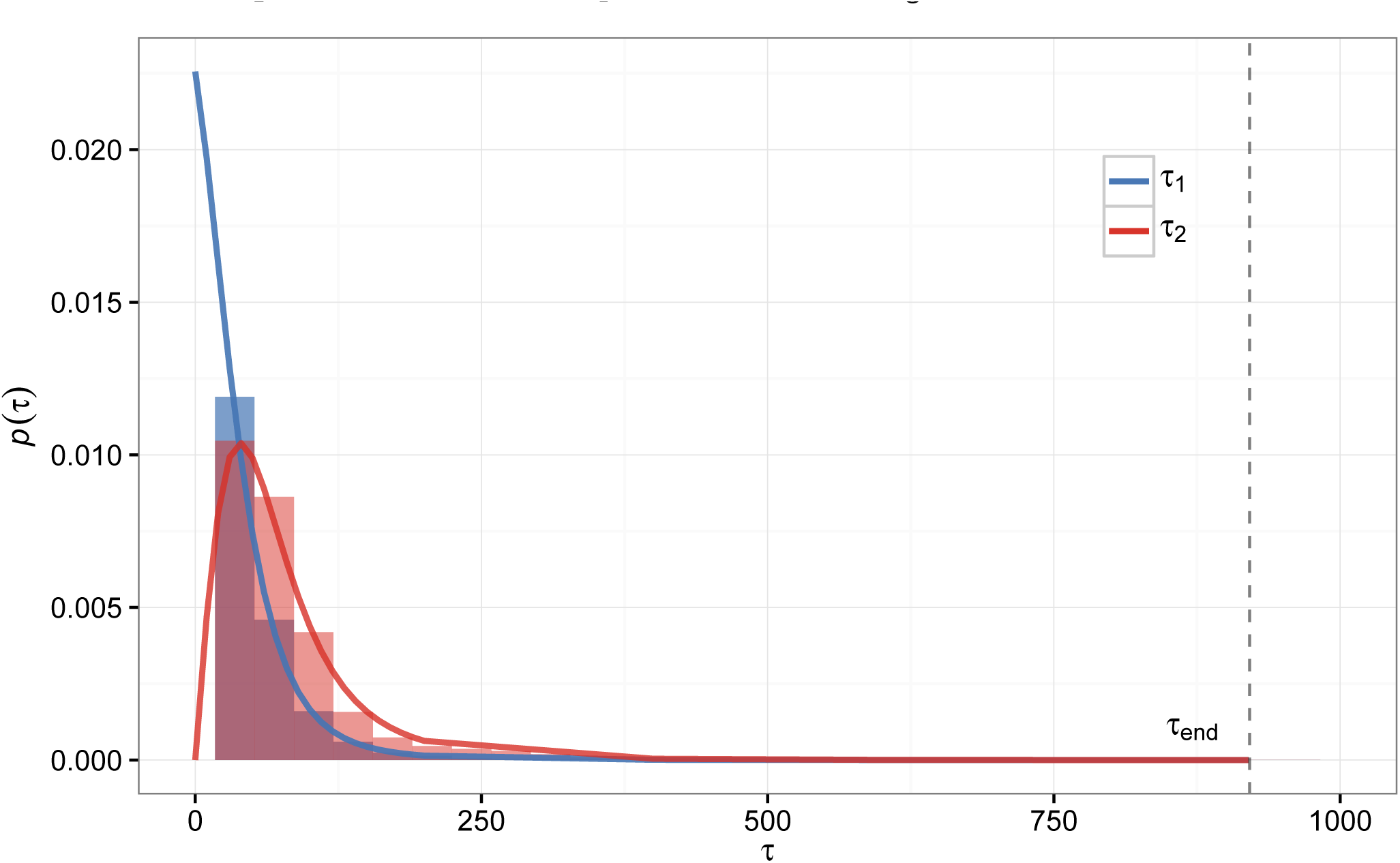
Waiting-time distributions for the establishment of the first and second mutants during rescue. Shown are the probability densities for the first and second mutants to establish during evolutionary rescue according to Equation (7) (blue line) and Equation (8) (red line). Empirical distributions from 10,000 simulations are shown in the same corresponding colors but as density histograms. The particular scenario is a low-density rescue scenario with *w*_0_ = 10,000, *K* = 110,000, *α* = 0.01,*b*_*m*_ = 1.1, and *w*_0_*μ* = 1.

### Soft sweeps are more likely when rescue is likely

Both *P*_rescue_ and *P*_soft_ vary similarly with the underlying parameters of our model because they both strongly depend on the area under the intensity function *R*(*t*). If we ask whether soft sweeps are more likely to be soft conditional on rescue occurring, we can see an obvious correlation between the two phenomena. This correlation is shown in Figure 6 where *P*_soft_|*P*_rescue_ is plotted against *P*_rescue_. Mathematically, we can derive the relationship using Equation (4) and Equation (6). Solving for *P*_soft_|*P*_rescue_ in terms of P_rescue_ gives

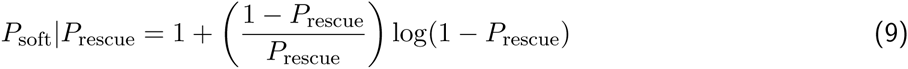

for *P*_rescue_ in (0, 1).

**Figure 6:**
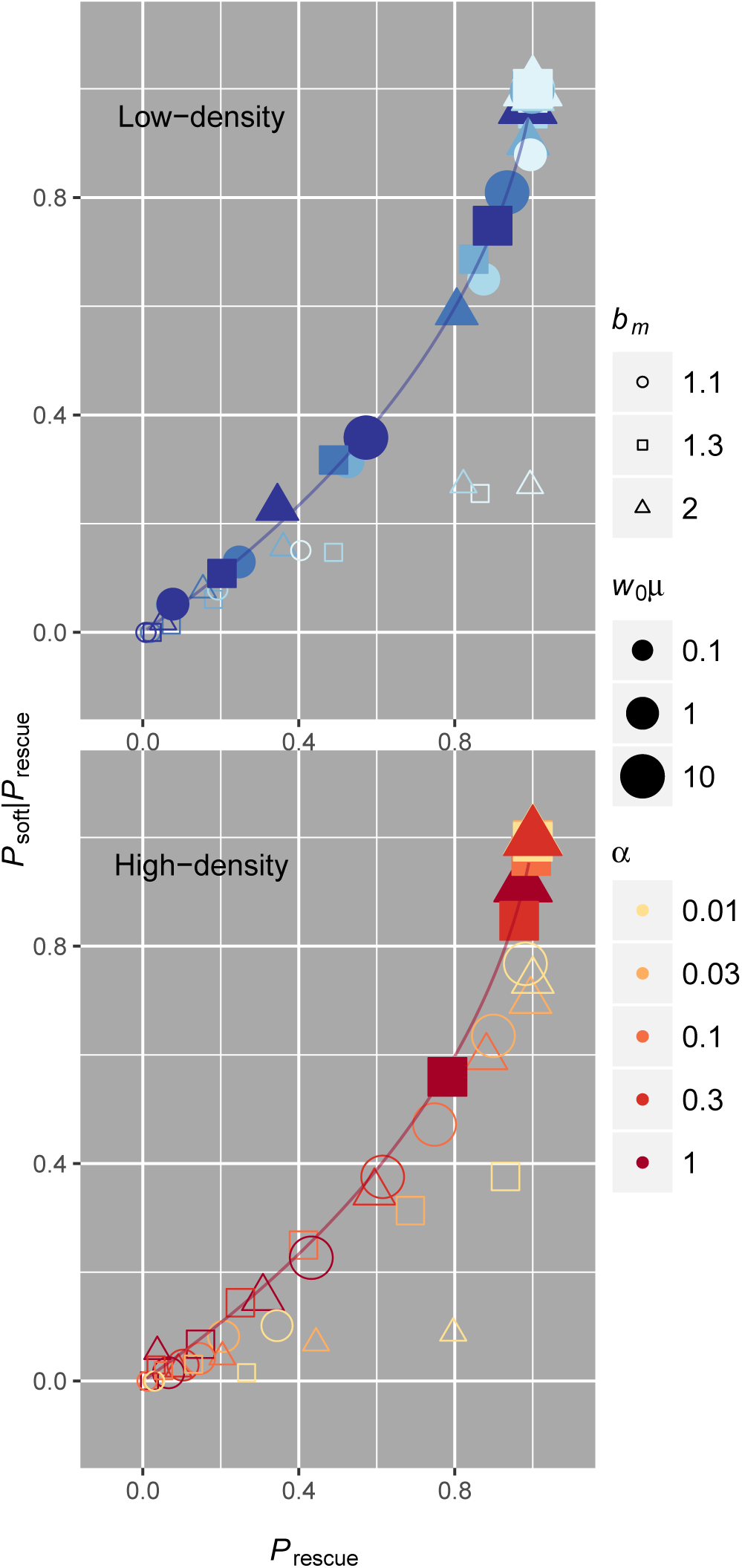
Frequent rescue is more likely to be driven by soft sweeps. Both in low-density (top, blue) and high-density (bottom, red) scenarios, simulations (points) indicate that soft sweeps are more prevalent when evolutionary rescue is likely. The correlation lines assume independence between lineages and are plotted according to the relationship in Equation (9). We have only shaded simulations where adaptation is not mutation-limited (*w*\**μ* ≥ 1, where *w*\* = *K*(1 – *d*_*m*_/*b*_*m*_)). When *w*\**μ* < 1, we expect our modeling assumptions to fail or soft sweeps to be unlikely. Note that the *w*\**μ* ≥ 1 criterion is met in all low-density simulations except where *w*_0_*μ* < 1, but departures where *w*\**μ* < 1 are more pervasive in high-density situations, especially when decline rates are slow.

While this correlation may be intuitive for many reasons, we highlight it because of its relevance to post-rescue genetic diversity. The hallmark of a soft selective sweep is that multiple lineages are preserved after selection (Hermisson and Pennings 2005; Pennings and Hermisson 2006a). This means that selection need not remove all genetic diversity in a population following evolutionary rescue, especially when rescue is expected to be common. We discuss why this might be important in the next section.

## Discussion

Understanding evolutionary rescue is important for characterizing population genetic aspects of environmental change, with applications ranging from conservation biology to preventing drug resistance evolution. Our results regarding the probability of evolutionary rescue agree with results previously published using similar models of adaptation under *de novo* mutation and adaptation from standing genetic variation (Orr and Unckless 2008; Uecker and Hermisson 2011; Martin *et al.* 2013; Orr and Unckless 2014). We further show that adaptation often proceeds via soft selective sweeps when evolutionary rescue is likely.

Our model captures important aspects of population dynamics and natural selection, although there are some limitations that we feel should be addressed. First, departures from our analytic assumptions occur in mutation-limited scenarios when population decline is very slow, namely establishment of one mutant lineage will affect the establishment of subsequent mutant lineages because of density-dependent mutant growth rates. Though we have chosen to not explicitly model this particular regime because it is not related to our primary focus on high recurrent mutation and soft sweeps, it is noteworthy to consider how populations can produce frequent rescue via hard sweeps when population decline is slow (illustrated in Figure 3, Figure 4, and Figure 6). Second, another drawback of our analysis is that our measure of *P*_soft_ is connected to the establishment of adaptive lineages during the process of evolutionary rescue (inferred from knowledge of the population composition after rescue or extinction) and not specifically connected to a sample genealogy, as in Pennings and Hermisson (2006a) and Wilson *et al.* (2014). This means that our measure of *P*_soft_ will not necessarily capture how the probability of observing a soft selective sweep depends on lineage frequencies. It is possible that lower frequency lineages could be missed in shallow samples. We can see how the observed relationship between average heterozygosity 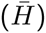 in our simulations (measured as the probability that all mutants sampled immediately following rescue are not identical by descent) and rescue probability has the same basic correlation as our measure of P_soft_|*P*_rescue_ in Figure 7. There is lower sensitivity to detect genetic diversity in such a shallow samples (sample size = 2 in Figure 7), but in larger samples, the expected correlation is virtually identical to the analytic expectation (sample size = 100 in Figure 7). We therefore highlight that empirical observations of soft selective sweeps in rescued populations are still crucially dependent sample depth.

**Figure 7:**
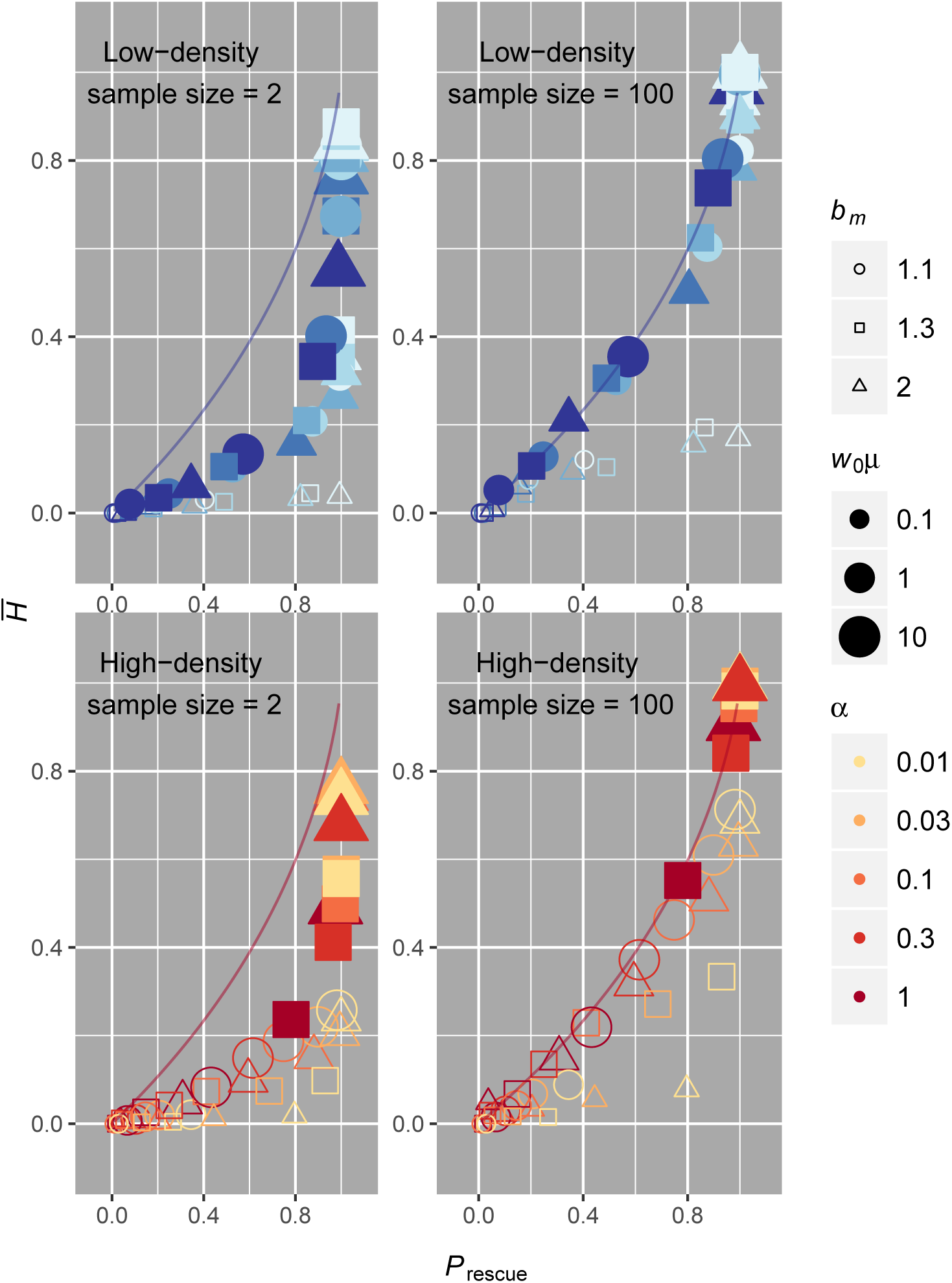
Mean heterozygosity from simulations of evolutionary rescue. Heterozygosity of the rescued population was calculated for each simulation and is equivalent to the probability that all sampled individuals do not come from a single mutant lineage. Plotted is the mean heterozygosity 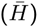 averaged over all 1,000 simulations for each parameter set against the corresponding probability of rescue for each parameter set. Only parameter sets where adaptation is not mutation-limited (*w*_0_*μ* ≥ 1 for low-density scenarios and *w*\**μ* ≥ 1 for high-density scenarios) are shaded. Although the sample depth is as low as possible for a meaningful measure of genetic diversity in the sample size equal to two case, we still see a general correlation between genetic diversity and probability of rescue. The sensitivity with which one could distinguish rescue probability using genetic diversity is much smaller compared to the previous measure of *P*_soft_ where we knew the precise number of lineages following evolutionary rescue. However, the sensitivity is virtually identical for larger samples, as seen in the sample size equal to 100 case. For larger sample sizes, the values of mean heterozygosity inferred from our simulations match our analytic predictions (colored lines) for situations where adaptation is not mutation-limited. This indicates that deeper population samples will have better sensitivity toward identifying whether rescue was likely or unlikely in scenarios where it is otherwise difficult to ascertain the probability of rescue.

### The generality of soft sweeps in frequent evolutionary rescue

We study soft selective sweeps via *de novo* mutation as a mode of evolutionary rescue because of their relevance to case studies of adaptation and particularly adaptation to strong environmental pressures (Karasov *et al.* 2010; Messer and Petrov 2013; Pennings *et al.* 2014; Feder *et al.* 2016). While adaptation from standing genetic variation is also expected to generate soft sweeps (Hermisson and Pennings 2005), it may be the case that adaptive mutants are absent at the onset of an environmental shift because they are strongly deleterious in the prior environment, as can be the case in resistance evolution (Andersson 2003; Shi *et al.* 2004; Cong *et al.* 2007). In reality, both modes of adaptation will play a role in the process of evolutionary rescue, and evolutionary rescue will depend strongly on the underlying ecological and population genetic factors of the adapting population such as population density, population substructure, epistasis, and genetic recombination. For example, whether fast decline of maladapted individuals in the population inhibits or facilitates evolutionary rescue depends strongly on whether adaptive mutations already exist in the population and whether there is strong population substructure (Wargo *et al.* 2007; Gatenby *et al.* 2009; Read *et al.* 2011; Uecker *et al.* 2014). And whether complex adaptations that require multiple mutations facilitate evolutionary rescue when a population faces an environmental challenge is strongly dependent on epistatic interactions between mutations and the presence (or absence) of genetic recombination (Lindsey *et al.* 2013; Uecker and Hermisson 2015).

Nevertheless, modeling evolutionary rescue as a Poisson process in each of these complex scenarios has led to a general form for to the probability of rescue as *P*_*rescue*_ = 1 – exp(–∧), where ∧ is the number of expected mutants generated via *de novo* mutation or existing in standing variation that are expected to survive extinction. While this number is a complex function of the aforementioned ecological and population genetic factors, the probability is nonetheless always higher when the number of surviving mutants is higher. Previous empirical observation (Bell and Gonzalez 2009) and theoretical intuition therefore suggest that when rescue is likely, more adaptive mutants are expected to be involved. It is for this reason that we expect soft sweeps to be a general feature of evolutionary rescue in situations where it is most likely to occur.

### The importance of genetic diversity in rescued populations

In Orr and Unckless (2014), the authors discuss how the average minimum population size that is reached during evolutionary rescue is smaller for adaptation via *de novo* mutation than for adaptation from standing genetic variation because of the dependence on the waiting time for the first established mutant ( *τ*_1_ in this paper). They posit that this may lead to lower genetic diversity in populations rescued via *de novo* mutation than those rescued from standing genetic variation, presumably because a larger average minimum population size could provide more opportunities for recombination to generate diversity within the rescued population. However, this relationship between genetic diversity and the mode of adaptation (from standing versus *de novo* variation) is not as direct when adaptation is driven by soft sweeps. Adaptation via soft sweeps may not drastically remove genetic diversity in an adapting population (Pennings and Hermisson 2006b), as might be expected in a hard sweep where only one lineage carries the adaptive mutation. In rescue via soft sweeps, adaptive mutations occur on different genetic backgrounds either before the environmental shift (in adaptation from standing genetic variation) or during the population decline after the environmental shift (in adaptation via *de novo* mutation). The degree to which each mode of adaptation reduces genetic variation will depend on the number of genetic backgrounds on which the adaptive mutation occurs. The number of different lineages could even be larger for adaptation via *de novo* mutation if the adaptive mutation is strongly deleterious (and correspondingly low in frequency) before the environmental shift, or it could be higher for adaptation from standing genetic variation if the adaptive mutation is already segregating on many different genetic backgrounds before the environmental shift. In either case, soft sweeps will play a significant role in preserving genetic diversity in the adapting population.

There are multiple reasons why higher genetic diversity following evolutionary rescue might be an important consideration. First, preserving genetic diversity that was present prior to the environmental shift will be important to the future fitness of the population following evolutionary rescue, especially in populations that cannot generate diversity quickly. If evolutionary rescue occurs via soft selective sweeps, then some of this ancestral diversity will be maintained for future generations. Second, post-rescue genetic diversity can be a useful proxy for measuring how likely such a population was to adapt to an environmental pressure. For example, post-treatment-failure genetic diversity could be used to determine the efficacy of a drug used to treat a virus within a patient in higher resolution than viral load alone because genetic diversity is expected to correlate with the likelihood that treatment failure occurred *a priori*. In other words, when treatment failure is common *P*_rescue_ is highest, and when *P*_rescue_ is highest we expect treatment failure to be driven by soft sweeps (Feder *et al.* 2016). This leads to higher genetic diversity in samples where failure was common and driven by soft sweeps than in samples where failure was rare and driven by hard sweeps. Indeed, others have found a correlation between treatment efficacy in HIV and whether treatment failure occurred via hard or soft sweeps that is in agreement with the theoretical results of this paper (Feder *et al.* 2016). Knowledge of this expected correlation may be broadly applicable to analysis of other types of drug-resistant infections, such as malaria. Ultimately, we believe that the basic characteristics of evolutionary rescue via soft sweeps described in our model will be useful for future studies.

## Methods

All code for the work performed in this article is available online at: https://github.com/benwilson87/evolutionary-rescue-soft-sweeps All plots were made using the ggplot2 (Wickham 2009) package in R (R Core Team 2015).

### Simulations

Simulations were performed using a Gillespie algorithm (Gillespie 1976) programmed in Python (Python Software Foundation. Python Language Reference, version 2.7.6, available at http://www.python.org). All simulations began at the onset of the wildtype population decline. The wildtype population was initialized at a population size of 10^4^ to be large enough to model the evolutionary processes by continuous approximations but small enough to be computationally efficient. For each round of simulation in the birth-death process, the algorithm used went as follows:

1. sample the waiting time *t* for an event from an exponential distribution with rate parameter equal to the sum of all rates for all possible events beginning at time 0,
2. randomly assign a specific event according to the relative probabilities of occurrence of each type of event (*i.e.* mutation, wildtype birth/death, mutant birth/death),
3. and update the population and all event rates for the new time *t*.

The process was repeated until either 1) the population went extinct and no rescue occurred or 2) adaptation successfully rescued the population and the new mutant population reached 99% of its equilibrium size. Note that in simulations where rescue occurred, we did not wait until the mutant subpopulation reached fixation. This was done to model the effects of sampling the population when rescue would likely be suspected rather than when complete replacement of the wildtype population occurred, a feature that we believe to be more realistic. At the end of each simulation, the population composition was analyzed to determine if 1) no lineages existed following an extinction event, 2) only one mutant lineage existed indicating a hard selective sweep, or 3) more than one lineage existed indicating a soft selective sweep. We simulated all combinations of:

1. five different wildtype decline rates: *α* ∈ {0.01,0.03,0.1,0.3,1.0},
2. three different mutation rates: *μ* ∈ {10^−5^, 10^−4^, 10^−3^},
3. three different mutant birth rates: *b*_*m*_ ∈ {1.1, 1.3,2.0}, and
4. two different population size limits: *K* ∈ {10000, 110000}.

We restricted birth rates to be relatively small to prevent bursting dynamics in the mutant population growth, however the absolute growth rates are potentially large at low population density. For all simulations, the wildtype and mutant death rates were set to 1. The population size limits were chosen to produce scenarios where the mutant would be unconditionally beneficial from the onset of the population decline (*K* = 110000) and where the mutant would initially suffer growth costs at high population density (*K* = 10000). Note that in the latter case, the mutant is actually deleterious with respect to the wildtype for all *α* except *α* = 1.0, in which case the mutant and wildtype are initially identical in fitness. The parameter ranges were chosen to cover a wide range of phenomenon with combinations that produced extinction almost always and combinations that produced rescue via soft selective sweeps almost always.

### Analysis

All mathematical analysis was numerically evaluated in Mathematica (Wolfram Research, Inc. 2010). In practice we found it neither necessary nor biologically meaningful to integrate to long times for all of our numerical integrations, so we chose to integrate to the time when a deterministically declining wildtype population would reach a single individual, *τ*_end_ = log(*w*_0_)/*α*, for our analytic calculations of *P*_rescue_ and *P*_soft_. For the waiting-time distributions for *τ*_1_ and *τ*_2_, we chose to integrate to the time when a deterministically declining wildtype population would reach 10% of its original size, *τ*_end_ = log(*w*_0_/1000)/*α*.

## Acknowledgments

We thank Hildegard Uecker for her comments and critique of the paper during its preparation. We thank members of the Petrov lab for comments and suggestions made prior to and during the formulation of this paper. B.A.W. was supported by the NSF Graduate Research Fellowship and NIH/NHGRI T32 HG000044. This work was also supported by the grant NIH 1RO1GM10036601 to D.A.P.

